# High-throughput selection of microalgae based on biomass accumulation rates in production environments using PicoShell Particles

**DOI:** 10.1101/2021.02.03.429271

**Authors:** Mark van Zee, Joseph de Rutte, Rose Rumyan, Cayden Williamson, Trevor Burnes, Randor Radakovits, Andrew Sonico Eugenio, Sara Badih, Dong-Hyun Lee, Maani Archang, Dino Di Carlo

## Abstract

Production of high-energy lipids by microalgae may provide a sustainable, renewable energy source that can help tackle climate change. However, microalgae engineered to produce more lipids usually grow slowly, leading to reduced overall yields. Unfortunately, tools that enable the selection of cells based on growth while maintaining high biomass production, such as well-plates, water-in-oil droplet emulsions, and nanowell arrays do not provide production-relevant environments that cells experience in scaled-up cultures (e.g. bioreactors or outdoor cultivation farms). As a result, strains that are developed in the lab often do not exhibit the same beneficial phenotypic behavior when transferred to industrial production. Here we introduce PicoShells, picoliter-scale porous hydrogel compartments, that can enable >100,000 individual cells to be compartmentalized, cultured in production-relevant environments, and selected based on growth and biomass accumulation traits using standard flow cytometers. PicoShells consist of a hollow inner cavity where cells are encapsulated, and a porous outer shell that allows for continuous solution exchange with the external environment so that nutrients, cell-communication factors, and cytotoxic cellular byproducts can transport freely in and out of the inner cavity. PicoShells can also be placed directly into shaking flasks, bioreactors, or other production-relevant environments. We experimentally demonstrate that *Chlorella* sp. and *Saccharomyces cerevisiae* grow to significantly larger colony sizes in PicoShells than in water-in-oil droplet emulsions (P < 0.05). We have also demonstrated that PicoShells containing faster biomass accumulating *Chlorella* clonal colonies can be selected using a fluorescence-activated cell sorter and re-grown. Using the PicoShell process, we select a *Chlorella* population that accumulates biomass 8% faster than does an un-selected population after a single selection cycle.

## Introduction

With the heightened interest in cell-derived bioproducts (e.g. high-energy lipids, recombinant proteins, antibody therapies, and nutraceuticals) and cell therapies (chimeric antigen receptor T-cell and stem cell therapies), the selection of desired cells based on their phenotypic properties has become increasingly important. In particular, the selection of microalgae strains for use as factories that convert light into biofuels has a long history because of their potential to be used as a carbon-neutral energy source. Specifically, high-energy lipids such as triacylglycerols (TAG) extracted from microalgae strains can be processed into biodiesel that can serve as an alternative energy source to power transportation^1,2^. CO_2_ emissions from the burning of biodiesel can later be fixed by microalgae and used to produce more high-energy lipids, creating a carbon-neutral mechanism to power today’s economy^3^. Microalgae are preferred over terrestrial plants because they have much faster biomass accumulation rates and particular strains won’t compete for resources that are important for agriculture^4^. Specifically, algae will occupy less land and certain strains can survive within recycled waste or seawater, eliminating potential competition for fresh water. In order to scale the microalgae industry to a point where microalgal-derived biofuels can be used ubiquitously, it is important to identify novel algae populations with enhanced biomass and lipid accumulation rates^5^. However, algal populations that are selected to overaccumulate high-energy lipids often have reduced growth rates^6^, making it necessary to develop technologies that can select algae populations based on their coupled biomass and lipid production.

Unfortunately, high-throughput screening tools for selection based on growth and bioproduct accumulation have not been readily available for scientists engineering cell strains. Fluorescence activated cell sorting (FACS) methods to select microalgal strains are only capable of selecting based on lipid content and not growth rate. This is because FACS traditionally has measured single cells at a single time point, rather than assaying colonies that are growing over time. Growth-based selection has been limited to low-throughput techniques such as using microtiter plates^7^ or small-scale bioreactors.

Microwell^8^, microcapillary^9^, droplet^10,11^, and gel microdrop technologies^12,13^ are capable of compartmentalizing single cells into nanoliter-sized compartments and allowing cells to grow into small clonal colonies for selection but do have some key limitations. The microfluidic approaches can have automated high-throughput selection mechanisms that make it possible to screen populations greater than 100,000 colonies or single cells per screen. Unfortunately, these microfluidic compartments have physical or chemical barriers that inhibit continuous solution exchange between the compartment and the external environment. In consequence, enclosed cells can rapidly deplete nutrients within the compartment, can accumulate secreted cytotoxic elements, change the pH of the environment, and cannot communicate with other cells via secreted factors. So, these high-throughput screening technologies may not be suitable for long-term (e.g. > 24 hour) growth assays, yielding selection pressures that are not aligned with final growth environments. This is because over these time scales, the compartments do not provide an environment that is physiologically similar to what is expected in scaled production cultures. As a result, selected cells may not behave the same way when scaled up for real-world applications as they did when cultured within these nanoliter-sized compartments. Scientists often need to perform further experiments and do additional genetic manipulation of selected strains to adapt them to scaled-up industrial cultures, a process that can take several months or years and without guaranteed success. Nanopen technology^14^ does have nanoliter-sized compartments that can have their solution replaced without dislodging the cells; however, it requires light-based manipulation of cells to isolate desired colonies that has a limited throughput of ~10,000 cells/screen.

To address these issues, we have developed a hollow shell microparticle platform (PicoShells), which enables compartmentalization of growing colonies, continuous media exchange, phenotypic screening and sorting via FACS, and viable downstream recovery. The PicoShell particles are ~90µm in diameter, consist of a solid outer shell made of polymerized (poly-ethylene glycol) PEG, and have a hollow inner cavity where microalgae can be encapsulated and cultured. More importantly, the solid PEG matrix is porous, allowing the PicoShells to be suspended in native media that is continuously exchanged, refreshing nutrients in the compartment and facilitating potential communication between cells in nearby compartments or in surrounding media. Additionally, cell-containing PicoShells can be placed directly into bioreactors or other relevant environments, providing a production-similar environment for enclosed cells that is not possible to attain with any other nanoliter-scale compartments. As a result of these features, strains developed using PicoShells are expected to exhibit their desired phenotypes in relevant scaled up cultures, promising to save cell-line developers months or years of additional labor to reach a similar point. These PicoShell particles can be loaded with single cells such that those cells grow over a multi-day period to form clonal colonies. Additionally, the pores in the outer hydrogel shell allow for encapsulated cells to be stained with common fluorescent tags such as BODIPY and live/dead stains. Since these particles are stable in aqueous solution, they can be screened and sorted using standard FACS instruments, potentially allowing colony-containing PicoShells to be sorted at throughputs >1000 particles/s. Cells can be released from PicoShells via mechanical or chemical mechanisms, retaining viability such that selected cells can be recultured, further scaled, analyzed, and perhaps re-sorted (Figure 1).

**Figure 1.**
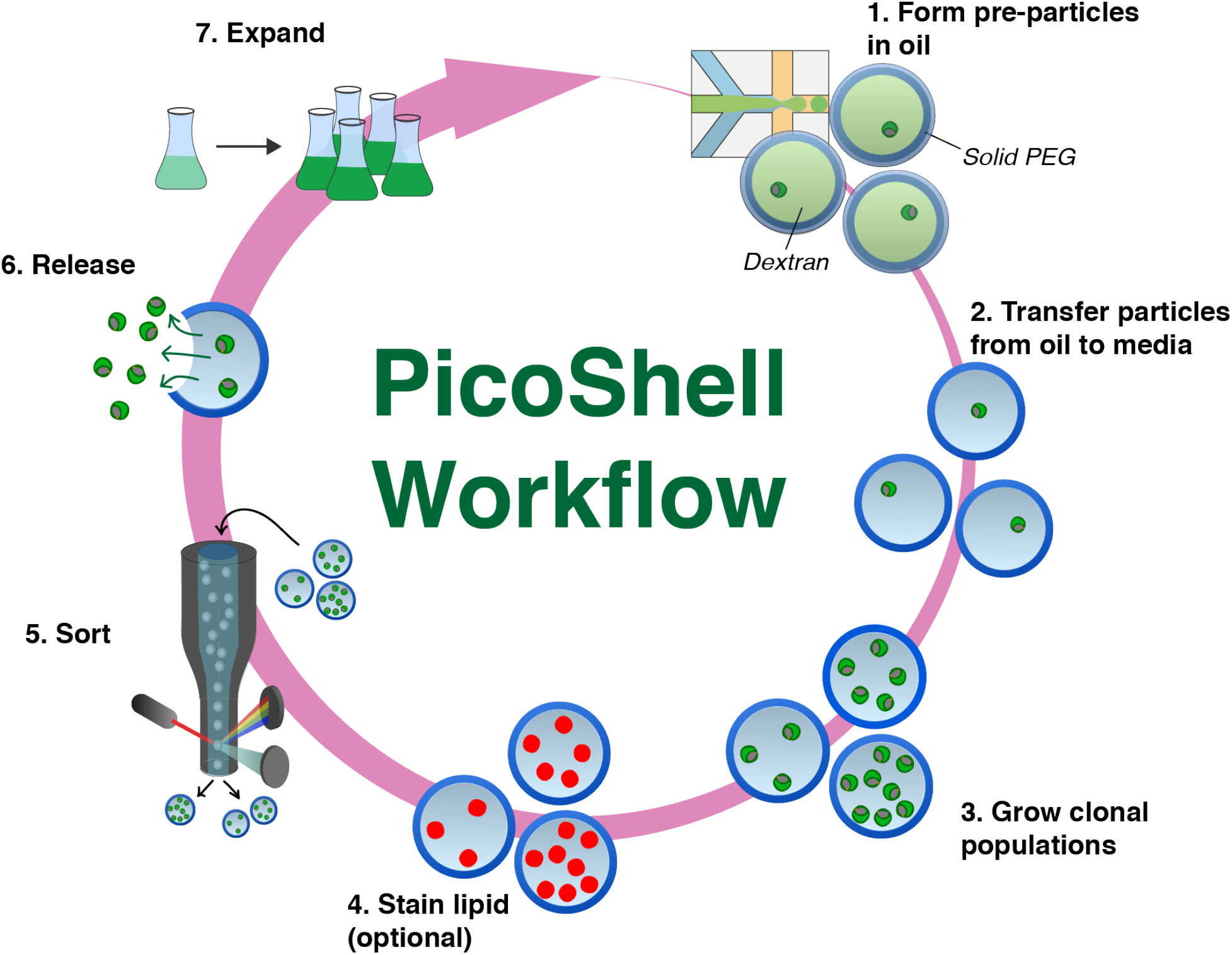
Workflow to enrich microalgae using PicoShell particles. (1) PicoShells are formed using droplet microfluidics, an aqueous-two phase system, and polymer chemistry. Particles are initially formed within an aqueous droplet surrounded by oil. Microalgae are within the dextran phase, which is surrounded by a solidifying PEG matrix. (2) Soon after particle formation, the particles are transferred into the algae’s native media. Pores in the solid outer shell allow for dextran to leak out and for continuous solution exchange. (3) Microalgae can grow within particles over multiple days to form clonal populations. (4) Pores in the solid matrix allow algal lipids to be fluorescently labeled. (5) High-performing populations can be sorted using FACS with scatter and/or fluorescence readouts. (6) Sorted particles can be broken down mechanically or by adding chemical reagents that degrade the PicoShell’s solid matrix, allowing associated cells to be released. (7) Released cells remain viable and can be re-cultured for further analysis and/or sorting.

Here, we compare growth of colonies in PicoShells with microfluidic droplet-in-oil approaches and demonstrate a proof-of-concept workflow for the selection of hyper-performing microalgae populations. In particular, we show how biomass accumulation of microalgae and yeast colonies is much faster in PicoShells than in water-in-oil droplet emulsions. Also, we demonstrate that particles containing faster growing algae colonies can be selected using FACS, that selected colonies can be released, and that the selected population can maintain a higher biomass accumulation rate than the un-selected population upon re-growth. We anticipate the PicoShell platform can play a key role in the selection of hyper-producing microalgae strains that translate to scaled-up culture environments as well as various other producer cell lines for a range of bio-products.

## Results

### Fabrication of PicoShells

PicoShell particles are made using a combination of microfluidic droplet technology^15,16,17^, aqueous two-phase systems (ATPS), and PEG polymer chemistry^18^ (Figure 2a). When mixed together at certain concentrations, a PEG-rich and dextran-rich phase can form, with a degree of phase separation that is tunable by adjusting the relative concentrations of the PEG and dextran components^19,20^ (Figure S1). Coalescence of the PEG-rich phase at different concentrations of PEG and dextran can lead to particles of unique shapes, owing to the unique interfacial tensions of the three-phase system (PEG-rich, dextran-rich, and oil phase)^21,22^. To determine the concentrations of PEG and dextran required to obtain PicoShell particles, we first obtained the binodal curve with the particular PEG and dextran used, a plot that defines the boundary between a completely mixed and phase separated aqueous two-phase solution. The binodal is dependent on the molecular weights and chemical functionality of the materials (Figure S2). We found that regions close to the binodal but above and to the right of the boundary led to the formation of concentric phases. When droplets contain PEG/dextran concentrations within 1-2% into the phase separation region above and to the right of the binodal, the dextran orients in the center of the droplet with PEG uniformly surrounding the dextran at the aqueous-oil interface (Movie 1). Crosslinking the PEG phase at these concentrations results in PEG hydrogel shells, i.e. PicoShell particles, that can remain stable when transferred out of oil and into aqueous solution (DPBS, media, etc.) (Figure 2b). The molecular weight of the dextran is chosen such that it can diffuse out of the enclosed shell particle following the phase transfer (Figure S3). The mechanism to form such hollow shell or capsule particles using the methods we describe has been adapted from previous work^23,24^.

**Figure 2.**
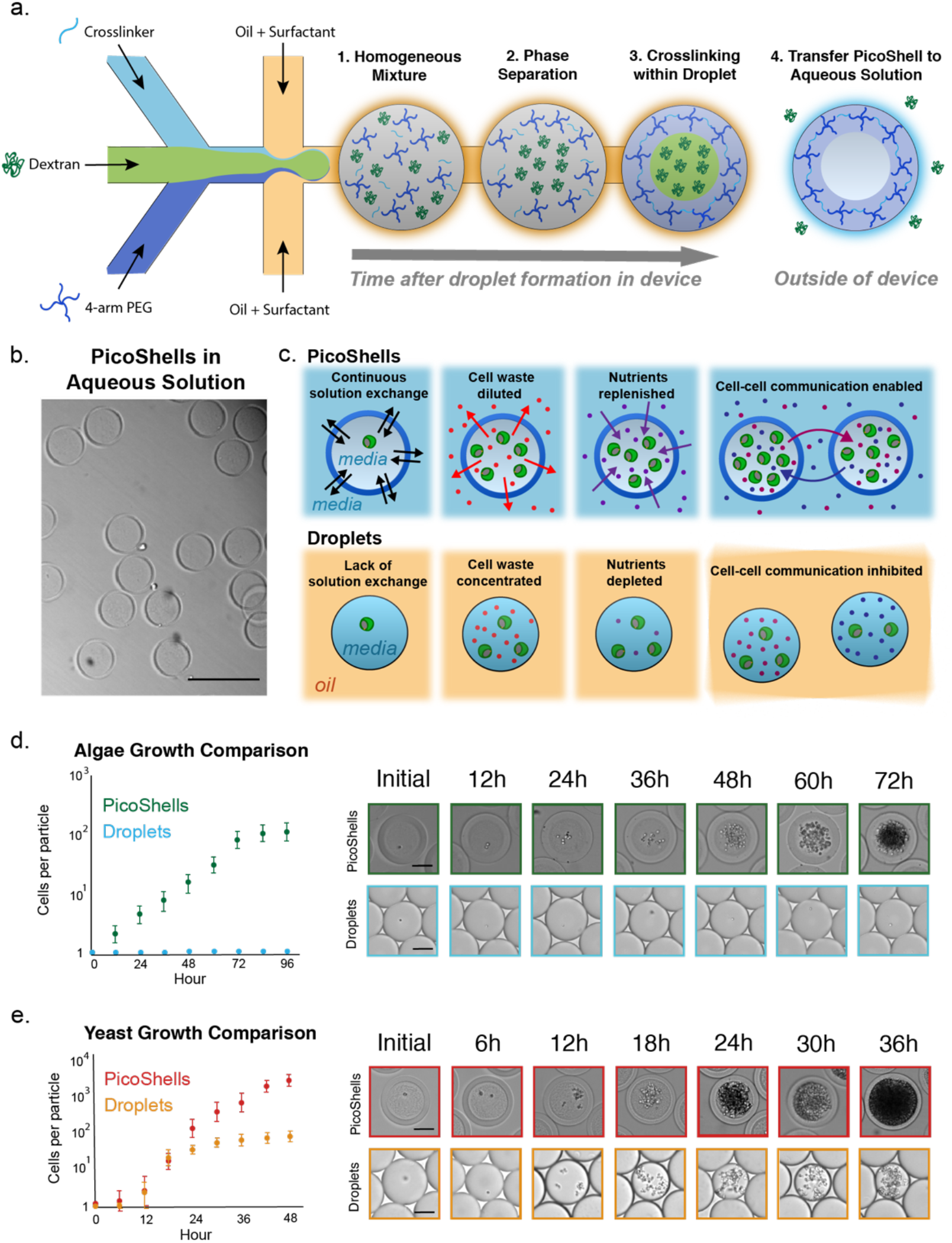
PicoShell particle formation and growth comparison between PicoShells and emulsion droplets. (a) PicoShells are formed by mixing together crosslinker, dextran, and 4-arm PEG to form a water-in-oil droplet emulsion. The reagents are mixed immediately before droplet formation to reduce premature gelation and phase separation. There is initially a homogeneous mixture of reagents, but 4-arm PEG and dextran phase separate as the droplet travels down the channel. As 4-arm PEG and dextran separate, the crosslinker and 4-arm PEG react. Following gelation, the particles are phase transferred from oil to aqueous solution and dextran leaks out of the particles via pores in the outer shell. (b) The co-flowing streams containing the three reagents can be distinguished in the device just prior to droplet formation. Particles naturally swell from 70µm to 80µm after phase transfer. Scale bars = 200µm. (c) *Chlorella* were encapsulated into PicoShells and droplets to compare growth rates in each compartment. Results show that the microalgae do not grow in droplets but grow readily in the particles. Scale bars = 50µm (d) *S. cerevisiae* were also encapsulated into PicoShells and droplets to compare growth rates. The yeast initially grow at the same rate in both compartments but growth eventually slows down in droplets. Scale bars = 50µm.

We identified a cell-compatible cross-linking chemistry to form PicoShell particles. While we can use UV-dependent chemistries to crosslink porous hydrogel particles of similar geometries^25^, we chose a different approach as UV-light and free radicals generated are likely to genetically or phenotypically affect cells being encapsulated, potentially introducing negative impacts on the assay/selection^26^. So, we incorporated a biocompatible pH-induced crosslinking chemistry where gelation occurs within physiologically-compatible pH ranges (pH ~6-8)^27^. However, pH-induced crosslinking introduces new challenges as the mixing time of crosslinker and functionalized PEG within droplets along with solidification affects the final particle morphology, even at the same PEG/dextran concentrations (Figure S4). If the crosslinking time is too quick, the PEG and dextran phases do not have enough time to fully phase separate, usually resulting in a non-uniform outer shell and/or un-desired crosslinking density in the cavity. In contrast, if the crosslinking time is too slow, the shift in the binodal resulting from the increasing molecular weight of the PEG phase as it starts to polymerize causes the formation of bowl-shaped particles instead. To obtain the ideal particle shape, we adjusted the crosslinking time by modulating the pH of the formed droplet. We found that repeatable, uniform shells could be formed by generating emulsions with in-droplet concentrations of 5% (w/w) 10kDa 4-arm PEG maleimide (PEG-MAL) crosslinked with dithiothreitol (DTT) and phase separated using 11% (w/w) 10kDa dextran. All reagents were dissolved at a pH of 6.25. With this combination of reagents, we are able to form uniform particles (Figure S5) with an outer diameter of 91 µm and shell thickness of 13 µm (CV of 1.7% and 6.9% respectively) at a particle generation rate of 720 PicoShells/s.

### Enhanced growth in PicoShells vs droplets

We found that encapsulated cells (*Chlorella* sp. and *Saccharomyces cerevisiae*) grow more rapidly and to higher final densities in PicoShells than in microfluidic droplets in oil (Figures 2c and 2d). Cells were from the same respective culture were encapsulated into PicoShells and droplets on the same day. *Chlorella* growth was tracked every 12h over a 72h period and *S. cerevisiae* was tracked every 6h over a 36h period. Interestingly, we found that *Chlorella* grow rapidly in PicoShells (Movie 2) starting with the formation of a first generation of daughter cells following 12h of incubation but did not grow when encapsulated in microfluidic droplets even over a 72h period (Figure 2c). *Chlorella* were encapsulated and incubated in autotrophic media, presumably making cells more susceptible gas transport. *Chlorella* were found to double every 12.2 hours and reached a carrying capacity within the 155 pL hollow cavity of a PicoShell of approximately 250 cells (Figure 2c). In parallel, we found that *Saccharomyces cerevisiae* grow both in PicoShells and in water-in-oil droplet emulsions (Figure 2d). However, while the growth rate of the yeast in both types of compartments were not statistically different before the first 18 hours of culture (P = 0.28 at 12h), the growth rate of droplet-encapsulated yeast became significantly slower than PicoShell-encapsulated yeast at later times (P < 0.001 at times > 18h). The reduction in the growth rate of droplet-encapsulated yeast is likely due to the depletion of essential nutrients and/or accumulation cytotoxic cellular waste. All nutrients present in both media types were below 200Da. Such nutrients are freely exchanged through the outer membrane of the PicoShells given the molecular weights below ~40kDa (Figure S3). This result is in agreement with the enhanced growth rate of *E*.*coli* observed when encapsulated in capsule particles compared with droplets in oil^24^. The average number of yeast cells in PicoShells is dramatically increased between 24 and 48 hrs after encapsulation to 2900 cells/PicoShell, ~20X higher than the carrying capacity in droplets (~150 cells/droplet).

Intriguingly, we also observed that *S*.*cerevisiae* do not stop dividing once they fully occupy the volume of the inner cavity of the particle and additional cells actually causes the particle to stretch, increasing the overall diameter (Figure S6). The diameter of the PicoShells can actually expand from an initial diameter of ~90µm to a maximum size of ~500µm after 4 days at which point the particle ruptures and releases the encapsulated cells (Movie 3). This phenomenon was not observed for encapsulated *Chlorella* colonies. Instead, the microalgae were observed to stop dividing when the colony reaches the carrying capacity of the particle.

### Sorting of Chlorella based on biomass accumulation rate

*Chlorella* colonies seeded and cultured in PicoShells were selected based on biomass accumulation rate using a FACS instrument. We used the colony’s chlorophyll autofluorescence, appearing in the Cy5 channel (ex:620, em:647), as a metric for biomass accumulation (Figure 3a). Generally, colonies containing greater numbers of cells also contain higher amounts of chlorophyll, generating higher Cy5 fluorescence readouts, as we have demonstrated in a previous study^12^. We demonstrated that lipids could also be stained through the PicoShells by mixing BODIPY with the colony-containing particles (Figure 3b). However, to simplify the study design to focus on improving the engineering aspects of the workflow, we only sorted clonal colonies based on biomass accumulation rate of chlorophyll and not lipid productivity. We encapsulated *Chlorella* at an average loading density, lambda of 0.1, which resulted in 91.7% of cell-containing particles with no more than a single cell.

**Figure 3.**
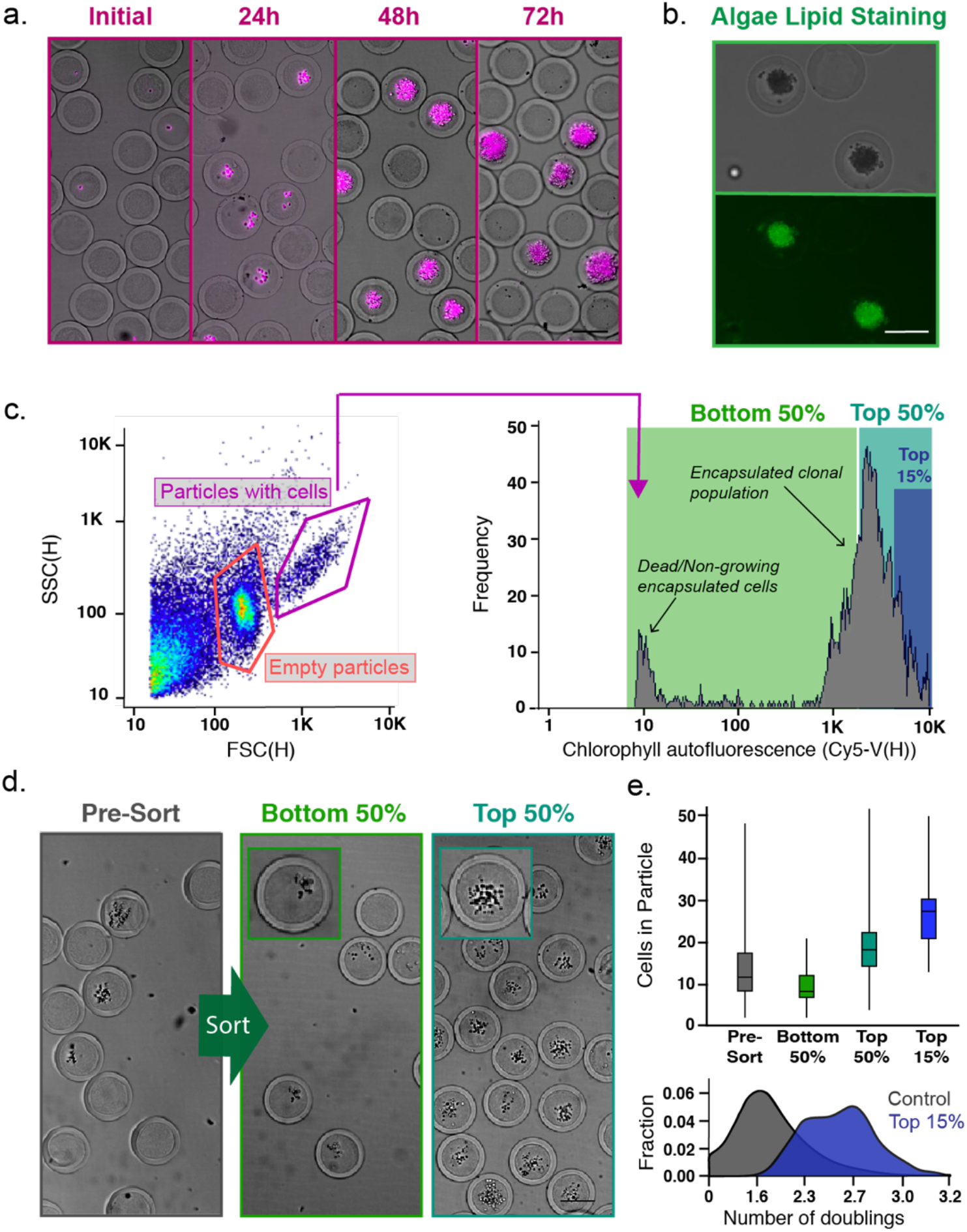
Screening and sorting characterization of microalgae-containing PicoShells. (a) PicoShells were loaded with *Chlorella* at lambda = 0.1 and allowed to grow for 48h. The biomass of *Chlorella* can be characterized via the chlorophyll autofluorescence that appears in the Cy5 channel. (b) The lipids in encapsulated *Chlorella* cells were stained with the addition of BODIPY 505/515. Localization of the stain were observed in the FITC channel. (c) After allowing *Chlorella* to accumulate biomass in PicoShells, the particles were screened using an On-Chip Biotechnologies Cell Sorter. Particles that contain colonies and cells can be distinguished from empty particles using scatter readouts. Colony-containing particles produce an observable Cy5 fluorescence distribution via the colony’s chlorophyll autofluorescence. (d) The colony-containing particles that produced the lowest 50%, highest 50%, and highest 15% Cy5 fluorescence readouts were sorted with 94.0% purity and 72.7% yield. 200 particles were sorted in each sample. (e) Selection of colony-containing PicoShells from different regions of the Cy5 distribution corresponds to particles containing different amounts of algal biomass, with particles with higher Cy5 fluorescence readouts containing more cells than those with lower Cy5 fluorescent readouts. Particles sorted from the higher end of the Cy5 distribution contain colonies that have undergone more doublings and have accumulated more biomass during the incubation period. Scale bars = 50µm.

Following culture for 48 hours, PicoShells containing *Chlorella* colonies were sorted using the On-Chip Sort at an average event rate of 100-200 events/sec. We observed three distinct populations in the forward scatter height [FSC(H)] vs side scatter height [SSC(H)] plot: one from colony-containing particles, one from empty particles, and one from debris (Figure 3c). The debris population was confirmed to be from particulates naturally present in *Chlorella* media. If the media is filtered, a greatly reduced fraction of debris events is observed (Figure S7). As expected, ~85.7% of detected particles do not contain cells due to the lambda = 0.1 that was used. In agreement with contrast observed in brightfield microscopy, PicoShells that contain microalgal colonies generally have increased forward and side scatter intensities. We verified that most of the colony-containing particles are within this high FSC/SSC gate by demonstrating that events in this gate also contained the highest Cy5 fluorescence (i.e. chlorophyll autofluorescence). A selected sample based on this gate had 94.0% purity of colony containing PicoShells (Figure S8).

Using the On-Chip Sort we selected out PicoShell particles with the fastest growing colonies by gating on chlorophyll autofluorescence. When we selected particles from different regions of the Cy5 distribution of colony-containing particles, we observed differing numbers of microalgae in the sorted colonies (Figure 3d). PicoShells gated on the lowest 50% in the Cy5 channel and within the high scatter gate possessed on average 9.2 ± 3.7 cells. This was statistically different from colonies recovered when gating the highest 50% (19.5 ± 7.1 cells, P < 0.0001) and highest 15% (27.0 ± 7.2 cells) (Figure 3e). Before sorting, colony-containing PicoShells contained on average 13.0 ± 7.7 cells. Overall, higher Cy5 fluorescence intensities corresponded to particles with a greater number of cells and given our loading conditions favoring single cell-derived colonies, it is likely these particles contained microalgae sub-populations that have faster doubling times and/or biomass accumulation rates (Figure 3e).

### Selection and re-growth of a hyper-performing Chlorella sub-population

We used the workflow to select *Chlorella* colonies based on Cy5 fluorescence, released colonies from the particles, re-cultured, and verified after re-culture that the selected sub-population accumulated biomass faster than an un-selected population (Figure 4a). For these studies we minimized PicoShells with more than one cell initially loaded by using lambda = 0.05, resulting in 3.2% of all particles containing colonies and ~98.3% of cell-loaded particles containing no more than one cell. Following 48h of growth in PicoShells immersed in *Chlorella* native media, we sorted colony-containing PicoShells gated to have the highest 11.1% of Cy5 fluorescence (425 events were selected from a population of 3839 colony-containing particles) (Figure S7).

**Figure 4.**
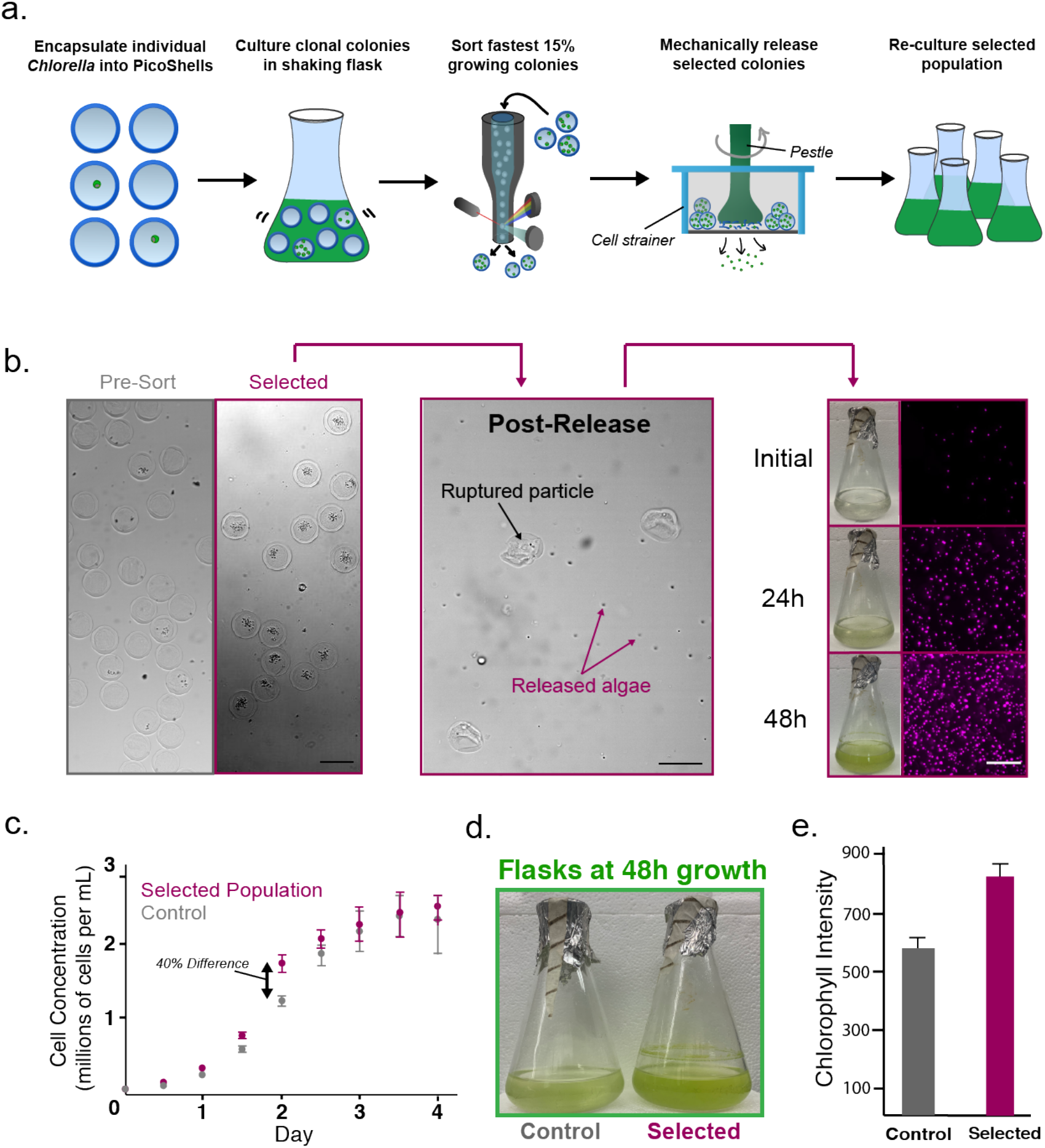
Selection of a hyperperforming *Chlorella* sub-population based on biomass accumulation rate (a) Single *Chlorella* were encapsulated into PicoShells and incubated under standard culturing conditions in a shaking flask to allow cells to accumulate biomass. Colony-containing PicoShells from the top 15% of the Cy5 fluorescence distribution were selected by FACS and mechanically released from particles. Released cells were then re-cultured for further analysis. (b) From a particle population of 121,213 particles (3839 containing colonies), 425 particles were selected. Selected particles were ruptured on top of a cell strainer, causing selected algae to be released into fresh culture media. This sample was re-grown in an Erlenmeyer flask under standard culturing conditions for several days. (c) The selected population and an un-selected population were seeded in separate flasks at the same concentration and their cell concentrations were tracked for 4 days. The selected population had an 8% faster growth rate (10.2h doubling times) than the un-selected population (11.2h doubling time) for the first 48h after seeding before slowing down as the culture reached carrying capacity. (d) The largest difference in biomass accumulation was observed at 48h after seeding (~40% difference in cell concentration), a difference that can be visibly seen in the green color of the cultures. (e) The difference in biomass accumulation was verified by measuring the chlorophyll density of each sample with a well plate reader at 48h after seeding. The selected population was measured to have a 27.6% higher chlorophyll density (P < 0.05). Scale bars = 100µm.

Selected colonies were released from PicoShells by applying mechanical shearing stress onto the particles, causing them to rupture (Movie 4). PicoShells were disrupted with mechanical shear and released cells were re-cultured in a flask (Figure 4b). We compared the biomass accumulation rate of the selected sub-population during re-culture to an un-selected population by seeding each population at the same concentrations and tracking their cell concentrations over a 4-day period (Figure 4c). We observed that the selected sub-population had an ~8% faster growth rate than the un-selected population (doubling times of 10.2h and 11.2h respectively, P < 0.01) for the first 48h of growth after seeding. This resulted in a 40% difference in the cell concentrations 48h after seeding that can be visibly seen in the culture flask (Figure 4d). The difference in accumulated biomass at this time point was also evaluated by measuring the total chlorophyll autofluorescence within well-mixed aliquots from each culture at this time point (Figure 4e), resulting in 27.6% increase in chlorophyll autofluorescence for the selected sub-population. As expected, the differences in cell number and overall chlorophyll biomass between the two populations diminished after 48h as the cultures reach carrying capacity.

## Discussion

### Advantages of PicoShells

Several key aspects of the PicoShell workflow suggest it can aid in the selection/evolution of cells and cell-based products including: (1) Cell behavior and growth is significantly enhanced in PicoShells compared to water-in-oil droplet emulsions. (2) PicoShells containing desired cells/colonies can be selected using commercial fluorescence activated cell sorters. (3) Selected cells/colonies can be successfully released from the PicoShells and re-cultured. (4) Selected populations maintained a high-growth phenotype post-process at least for several generations. Importantly, PicoShells can be placed and remain stable in more production-relevant environments (e.g. a shaking culture flask, bioreactor, outdoor cultivation farms) that are not possible with other high-throughput selection technologies (e.g. droplet technology, microwells, etc). The porous outer shell enables solution exchange with the external environment, allowing replenishment of nutrients, diffusive transport and dilution of cytotoxic cellular waste, access to quorum sensing factors from external cells/colonies, and exposure to natural concentration, temperature, light, or physical gradients in the culture environment. As a result, PicoShell technology may provide a novel high-throughput screening tool that enables cell line developers and researchers to select cells based on their behavior in production-relevant environments, making it much more likely that selected populations will exhibit the desired phenotypic properties when scaled up for real-world applications.

Growth of other algae species in droplets has been previously shown^11, 12, 30, 31^, making it intriguing why *Chlorella* in particular does not survive when encapsulated into water-in-oil droplet emulsions. While it is unclear exactly why this particular phenomenon occurs, we believe that the lack of cell survival is related to the restricted gas exchange across the oil barrier. This particular species is grown in autotrophic media and is very sensitive to gaseous CO_2_ concentrations. We have observed that bulk cultures of this particular species cannot grow when not cultured in an incubator that regulates CO_2_ or not cultured with media that is not supplemented with sodium bicarbonate^32^. While previous studies have shown that gases can generally pass through fluorinated oil^33, 34^, this diffusion may be limited or altered to an extent that sensitive species are greatly affected unlike more robust cell types (*Chlamydomonas reinhardti, Euglena gracilis*, etc). Regardless of the root cause for the lack of growth in droplets, the results demonstrate that the environments in PicoShells and droplets are different enough that we can observe a noticeable effect on cell behavior, a result that is substantiated by the improved growth properties of *S. cerevisiae* in PicoShells.

### Potential for chemically degradable PicoShells

We have explored multiple mechanisms to chemically release cells from PicoShells by including chemically-degradable motifs in the outer PEG shell. Currently, we can consistently fabricate PicoShells crosslinked with PEG-MAL and DTT. These are compatible with multiple cell types, including *Chlorella, S. cerevisiae*, and *Euglena gracilis* (Figure S9). PicoShells crosslinked with PEG-MAL and DTT can be broken down with the addition of sodium periodate (NaIO_4_) due to presence of a diol in DTT. Unfortunately, NaIO_4_ can be toxic^28^ and likely kills or has large negative impacts on many cell types. Previously, we have made hydrogel particles with degradable peptide crosslinkers^18^, and similar incorporation of degradable crosslinkers could enable enzymatic or chemical degradation of particles to release selected cells/colonies. As an initial proof of concept of this approach, we developed PicoShells that contain di-sulfide linkages that can be degraded via the addition of DTT or TCEP. *S. cerevisiae* encapsulated in these particles, remain viable, grow, and can be chemically released (Figure S10 and Movie 5). Unfortunately, a chemical precursor we use to form these particles (4-arm PEG-Ortho-Pyridyldisulfide) is toxic to *Chlorella* (Figure S11), suggesting that the chemical formulation of the PicoShell should be matched to the cell type. We have also encapsulated and grown *Chlamydomonas reinhardtii* in PicoShells crosslinked with matrix-metalloproteinease (MMP)-degradable-peptides (Figure S12) that can be degraded with the addition of trypsin (Movie 6). Unfortunately, *C. reinhardtii* (and likely other cell types) naturally secrete MMPs that often pre-maturely break down the particles^29^.

While the mechanical mechanism of release we demonstrate works well for releasing bulk populations of selected particles, it is likely difficult to adapt the process to separately release individual colonies (e.g. a single particle in a single well). Such single particle isolation is important if a researcher wishes to explore the different strategies for hyper-performance and the various underlying genetic mechanisms that result in such phenotypes. While it may be possible to engineer tools to mechanically break down a single particle, release of cells using these tools may be complicated and inefficient. Hence, it may be necessary to fully develop PicoShells that are chemically degradable and compatible with several cell types. While we have engineered disulfide crosslinked PicoShells that are compatible for yeast applications, we have also shown that it is difficult to discover chemistries that enable chemical degradation and maintain cell viability for more sensitive cell types.

### Limitations on throughput

We have also found that there is a tendency for crosslinked material to stick to the walls near the droplet generation junction, causing a disruption in the flow (Figure S13). Since we use pH-induced gelation and the gellable materials (PEG-MAL and DTT) come into close proximity briefly before droplet generation, gelled material often forms at the junction, inducing jetting and disruption of particle formation approximately 15 minutes after initial particle formation. As a result, the device needs to be replaced each time particle formation is halted, reducing the overall number of PicoShells that can be manufactured to 370,000 particles per device.

The jetting of reagents due to pre-mature formation of gelled material that sticks to the walls of the droplet generator limits the overall throughput of PicoShell generation. Use of UV-induced crosslinking mechanisms can solve this problem since gelation would occur downstream of droplet generation^27. 35, 36^, unlike pH-induced mechanisms where mixing of reagents immediately prior to droplet generation often results in gelled material forming in the droplet-generation junction over time that disrupts the overall flow. However, use of UV-induced crosslinking likely creates issues for particular cellular applications, as previously discussed. At the same time, UV-induced crosslinking may be used for workflows involving resilient cell types (e.g. bacteria) or workflows where cells are mutagenized prior to selection, and UV-induced mutations would be potentially beneficial. In another approach, we can use a 3D-printed device that can fabricate PicoShells and is composed of a material that reduces the amount of gelled material that sticks to the device walls^37^. A summary of the different types of PicoShells that can we can currently fabricate and their advantages and disadvantages is shown in Table 1.

**Table 1.**
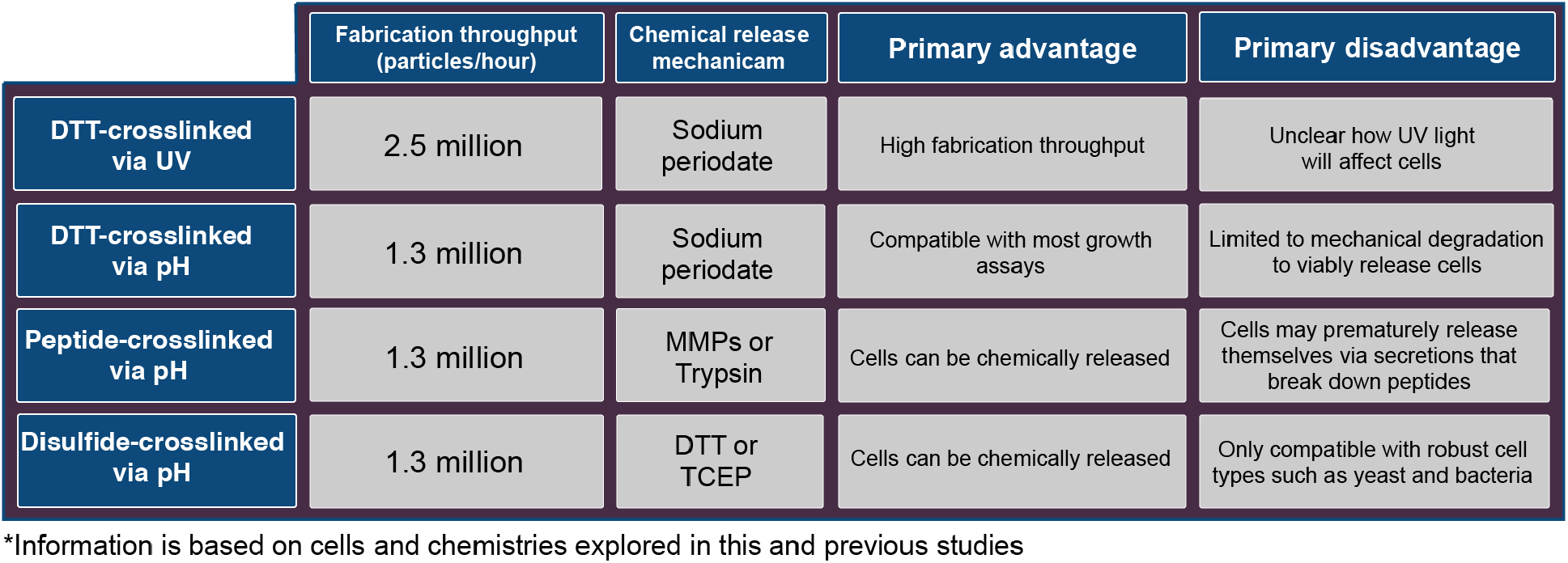
Summary of Current PicoShell Variations.

### Potential future applications

Despite these solvable limitations, the experimental evidence we have presented shows that PicoShell technology has significant advantages. The workflow can be potentially used for directed evolution of cell populations^38^ where mutagenized cells are placed under selection pressures to generate novel strains, based on unique selection criteria that are time-dependent (e.g. growth / biomass accumulation), at the colony level (multi-cellular construct formation), or require solution exchange steps (lipid staining, ELISA-based assays). For example, the technology may be used to produce microalgae strains that overperform in lipid accumulation rates without significantly reducing their rate of biomass accumulation for biofuel applications. The technology may also be used to generate novel yeast strains that maintain high biomass production at higher ethanol concentrations, potentially enhancing the overall production of ethanol biofuels^39,40^, plastics^41^, materials^42^, and alcoholic beverages^43^.

The outer shell’s PEG material is also able to be modified, enabling the technology to be potentially used for relevant mammalian cell applications. For example, affinity motifs such as antibodies and peptides can be added to the solid matrix that can capture cellular secretions^27^. Antibody-conjugated PicoShells may be used to produce hyper-secreting and hyper-growing CHO cell populations based on their behavior in bioreactors for scaled production of protein therapeutics. The pore size of the particles may also be modulated by changing the MW of PEG used to crosslink the solid phase^45^ or by including non-functionalized PEG^46^, gelatin^47^, or hyaluronic acid^48^ in the PEG phase. Adherence motifs such as RGD peptides, fibronectin, or poly-L-lysine (PLL) may be also added to the outer PEG matrix so that stem cells, adherent CHO cells, or other adherent cell types have a solid surface to adhere to, further expanding the potential applications of the PicoShell workflow.

In summary, we have shown that PicoShells may enable cell line developers to develop novel cell populations based on their behavior in production environments. Unlike previously developed high-throughput screening tools, individual cells can be compartmentalized, placed into relevant environments such as bioreactors, exposed to natural stimuli, and selected based on their time-dependent behavior and growth in such environments via widely used flow cytometers. As a result, the technology has the exciting potential to rapidly accelerate the development of cell-derived bioproducts such as biodiesel, materials, cell-derived agriculture, nutrient supplements, and protein therapeutics.

## Materials and Methods

### Bulk culture of cells

*Chlorella* cells (CCMP1124 from National Center for Marine Algae and Microbiota) used in the study were provided by Synthetic Genomics, Inc. *Chlorella* populations were cultured in 500mL Erlenmeyer flasks containing seawater based medium with added vitamins, trace metals, nitrate, phosphate, and sodium bicarbonate (SM-NO3 medium)^49^. SM-NO3 medium was also supplemented with penicillin-streptomycin (P/S, Thermo Fisher Scientific, 15140122). *Chlamydomonas reinhardii* (STR CC-4568) and *Euglena gracilis Z* (NIES-48) procured from Microbial Culture Collection at National Institute for Environmental Studies (NIES) Japan were cultured in 500mL using Tris-acetate-phosphate medium^41^ and Koren-Hunter (KH) medium at a pH of 5.5^50^ respectively. Flasks containing algae cultures were shaken continuously at 120RPM with constant 150µE light at room temperature. Algae cultures were kept at a concentration of 2-10 million cells/mL. Strains of *Saccharomyces Cerevisiae* were obtained from Sigma-Aldrich (STR YSC1). The yeasts were grown in yeast extract (1%, w/v) peptone (2%, w/v) glucose (2%, w/v) (YPD) media supplemented with 50mg/L ampicillin (Sigma-Aldrich, 69534). The strains were grown in 250 mL Erlenmeyer flasks containing 100 mL of YPD, under aerobic conditions at 30°C with agitation (300 rpm).

### PicoShell Fabrication

Mechanically-degradable particles demonstrated throughout the majority of the study were fabricated forming uniform water-in-oil droplet emulsions containing in-droplet concentrations of 5% (w/w) 10kDa 4-arm PEG-maleimide (4-arm PEG-MAL, Laysan Bio), 11% (w/w) 10kDa dextran (Sigma Aldrich, D9260), and 1.54mg/mL dithiothreitol (DTT, Sigma Aldrich, 10197777001). Reagents were dissolved into SM-NO3 medium, YB, TAP, or KH medium for the encapsulation of *Chlorella, S. cerevisiae, C. reinhardtii*, or *E. gracilis* respectively (each at a pH of 6.25). Novec™ 7500 Engineeried fluid (3M™, 297730-92-9) with 0.5% Pico-Surf™ (Sphere Fluidics, C024) acting as surfactant was used as the continuous, oil phase. Droplet emulsions were formed using a 4-inlet microfluidic channel fabricated with polydimethylsiloxane (PDMS) using standard soft-lithography techniques^51^. Reagents were loaded into separate syringes and pushed through the PDMS droplet generator using syringe pumps (Harvard Apparatus, MA, USA). In order to reduce the effects of functionalized PEG and crosslinker on cell growth during encapsulation in PicoShells, cells were suspended in the dextran phase such that the cells only interact with the PEG and DTT reagents for a short period of time (Movie 7). In-droplet concentrations of 3.25% (w/w) 20kDa 4-arm PEG Ortho-Pyridyldisulfide (4-arm PEG-OPSS, Creative PEGWorks, PSB-459), 10% (w/w) 10kDa dextran, and 0.80mg/mL DTT were used to form di-sulfide linked PicoShells. In-droplet concentrations of 5% (w/w) 10kDa 4-arm PEG-MAL, 11% (w/w) 10kDa dextran, and 14.1mg/mL di-cysteine modified Matrix Metallo-protease (MMP) (Ac-GCRDGPQGIWGQDDRCG-NH_2_) (Genscript) peptide substrate were used to form MMP-degradable PicoShells.

Following droplet generation, emulsions were stored at room temperature for 1h to allow PicoShells to fully solidify. The PicoShells were de-emulsified by adding Pico-Break™ (Sphere Fluidics, C081) at a 1:1 volume ratio on top of the PicoShells. Once Pico-Break had passed through all the PicoShells, the particles were transferred into aqueous solution (DPBS or cell media) containing 10µM N-ethylmaleimide (NEM, Sigma-Aldrich, E3876) at a pH of 6.5. The PicoShells were kept in NEM solution for 0.5h to allow NEM to react to any free thiols on the particles to reduce clumping. PicoShells were then passed through a 100µM cell strainer to remove any oversized or clumped particles and a 40µM cell strainer to remove any free cells or small debris before being transferred into cell media to be used for the particular assay.

### PicoShell versus droplet emulsion growth comparison

*Chlorella* and *S. cerevisiae* from the same respective initial culture were separately encapsulated into mechanically-degradable PicoShells and microfluidically-generated droplets in oil of approximately the same volume (155pL) using the same droplet generator. Each cell type was encapsulated into PicoShells and droplets within 2h of each other. PicoShells and droplets containing *Chlorella* were incubated in Eppendorf tubes with constant 150µE light at room temperature (no shaking). Compartments containing *S. cerevisiae* were incubated in Eppendorf tubes at 30°C (no shaking). Both PicoShells and droplets were not shaken since droplets tend to de-emulsify when shaken at speeds >100RPM. PicoShells and droplets were imaged using an inverted microscope in BF and Cy5 (ex: 620nm, em: 676nm) fluorescence at equal time intervals over a multi-day period to track the growth of cells in their respective compartments over time.

### Staining of Intracellular Lipids

Following a 48h culture of *Chlorella* in PicoShells, intracellular lipids were stained with BODIPY^505/515^. Stock BODIPY^505/515^ was prepared by dissolving 4,-Difluoro-1,3,5,7-Tetramethyl-4-Bora-3a,4a-Diaza-s-Indacene (Life Technologies, D3921) powder into dimethyl sulfoxide (DMSO) at a concentration of 2.5mg/mL and then diluted to 2.5µg/mL using SM-NO3 media. Colony-containing PicoShells were placed at a concentration of 2 x 10^6^ particles/mL in SM-NO3 media before being mixed at a volume ratio of 1:1 with 2.5µg/mL BODIPY^505/515^ and incubated in the dark for 0.5h. The PicoShells were washed three times with SM-NO3 before being imaged in the FITC channel (ex: 488nm /em: 543nm) using a fluorescence microscope.

### Incubation and flow cytometric sorting of PicoShells

*Chlorella* were encapsulated into 90µm diameter PicoShells following Poisson loading with lambda = 0.1 for the initial sort and lambda = 0.05 for the full selection and placed into SM-NO3 medium at a particle to media volume ratio of 1:50. The particle-containing solution was then placed in a 250mL Erlenmeyer flask shaking at 120RPM and at room temperature under constant 150µE light for 48h to allow cells to accumulate biomass.

Colony-containing PicoShells were screened and sorted using an On Chip Sort (On Chip Biotechnologies, USA). The cytometer was equipped with both 488nm and 561nm excitation lasers and a PE-Cy5 (676/37nm) filter. Events were triggered based on particle absorbance from the 488nm laser. PicoShells were sorted based on their scatter readouts and thresholding desired intensity heights through the PE-Cy5 filter. PicoShell solutions were concentrated in fresh SM-NO3 media at a 1:10 particle to media volume ratio for screening and sorting. PicoShells within the selection gates were dispensed in a single collection reservoir. The sorted particles were then imaged using an inverted microscope and the number of cells in each particle was counted using MATLAB code.

### Release of cells and re-culture of selected populations

Post-selection, *Chlorella-*containing PicoShells were placed onto a 37µm cell strainer and placed over a 15mL conical tube containing fresh SM-NO3 media supplemented with P/S. The PicoShells were then ruptured by ‘grinding’ the particles with a pestle and washing with SM-NO3 media for ~5min, causing released cells to fall through the pores of the cell strainer and into the fresh media. Despite being able to be ruptured by direct mechanical shearing pressure, PicoShells remain stable in adverse indirect mechanical shearing pressures such as mixing, vigorous pipetting, and vortexing. The solution containing released cells was then transferred into a 250mL Erlenmeyer flask and put in standard bulk *Chlorella* culture conditions for 7 days to allow released cells regrow to a concentration of 15-20 million cells/mL.

To test for maintenance of an enhanced biomass accumulation phenotypes in the selected population, we seeded the selected population and an un-selected population into separate 250mL Erlenmeyer flasks with SM-NO3 media supplemented with P/S at a concentration of 500,000 cells/mL. The flasks were placed side-by-side under standard *Chlorella* culturing conditions for 4 days. The cell concentration was measured using a hemocytometer every 12h. At 48h of growth, we also measured biomass accumulation by aliquoting several fractions from the selected and un-selected sample into a well plate and measured the chlorophyll density (parameters) using a well plate reader at this time point.

### Chemically-induced degradation of PicoShells

To chemically degrade the various types of PicoShells, we first diluted or concentrated PicoShells to a concentration of 1 x 10^6^ particles/mL and added the following reagents at the indicated final concentration to degrade each PicoShell type: 10µg/mL sodium periodate (NaIO_4_, Fisher Scientific, P120504) for particles crosslinked with 4-arm PEG-MAL and DTT, 10mg/mL DTT or 3mg/mL Tris(2-carboxyethyl)phosphine (TCEP, Sigma-Aldrich, 646547) for particles crosslinked with 4-arm PEG-OPSS and DTT, and 0.0025% Trypsin with EDTA (Thermo Fisher Scientific, 25300120) for particles crosslinked with 4-arm PEG-MAL and di-cysteine modified MMP degradable peptide.

## Acknowledgements

This work was supported by the Presidential Early Career Award for Scientists and Engineers (N00014-16-1-2997) and California NanoSystems Institute at UCLA Planning Award.

## Author contributions

J.D., M.v.Z., and D.D conceived and designed initial PicoShell systems and protocols. J.D. designed the fabrication workflow for PicoShells and performed dextran diffusion experiments. J.D., M.v.Z., and M.A fabricated PicoShells. M.v.Z. and R.Ru. developed algae biomass assay protocols, performed experiments, and analyzed data. M.v.Z. and C.W. developed yeast biomass assay protocols, performed experiments, and analyzed data. T.B. and M.v.Z. performed algae growth comparison assays and analyzed data. A.S., D.L. S.B., and M.v.Z. performed yeast growth comparison assays and analyzed data. M.v.Z. and R.Ru. designed and performed flow sorting studies. M.v.Z. designed and performed the full *Chlorella* selection assay and analyzed data. R.Ra. provided valuable microalgae insights and expertise. M.v.Z. and D.D. wrote the manuscript with inputs from all authors. M.v.Z prepared figures. D.D. supervised the study.

