## Supplemental Information for "High-throughput selection of microalgae based on biomass accumulation rates in production environments using PicoShell Particles"

**Movie 1.** ATPS and PicoShell formation in droplet generator. PEG and dextran concentrations are fine-tuned such that a dextran-rich phase forms in the center of the droplet and a PEG-rich phase forms at the droplet's aqueous-oil interface. Crosslinking begins while the droplet travels downstream of the location of droplet formation.

**Movie 2.** Growth of *Chlorella* in PicoShells. Noticeable division of *Chlorella* in PicoShells is observed over a 24-hour period.

**Movie 3.** PicoShell rupture due to *S. cerevisiae* biomass accumulation. After ~80 hours of *S. cerevisiae* growth following initial encapsulation, the yeast cells can accumulate sufficient biomass such that the PicoShell diameter stretches from ~90µm to ~500µm before rupturing.

**Movie 4.** Mechanical breakdown of PicoShells. *Chlorella*-containing PicoShells were compressed between a glass slide and cover slip. The cover slip was pressed down onto the particles and twisted to cause PicoShells to break apart and cells to be released.

**Movie 5.** Chemical breakdown of disulfide-linked PicoShells. PicoShells containing disulfide linkages can be dissolved with the addition of DTT.

**Movie 6.** Trypsin breakdown of MMP-degradable-peptide-linked PicoShells. PicoShells crosslinked with peptides can be broken down with the addition of proteases such as trypsin.

**Movie 7.** Encapsulation of *Chlorella* into PicoShells. Cells are placed into the dextran phase prior to PicoShell fabrication. *Chlorella* can be visualized in the Cy5 fluorescence channel during encapsulation.

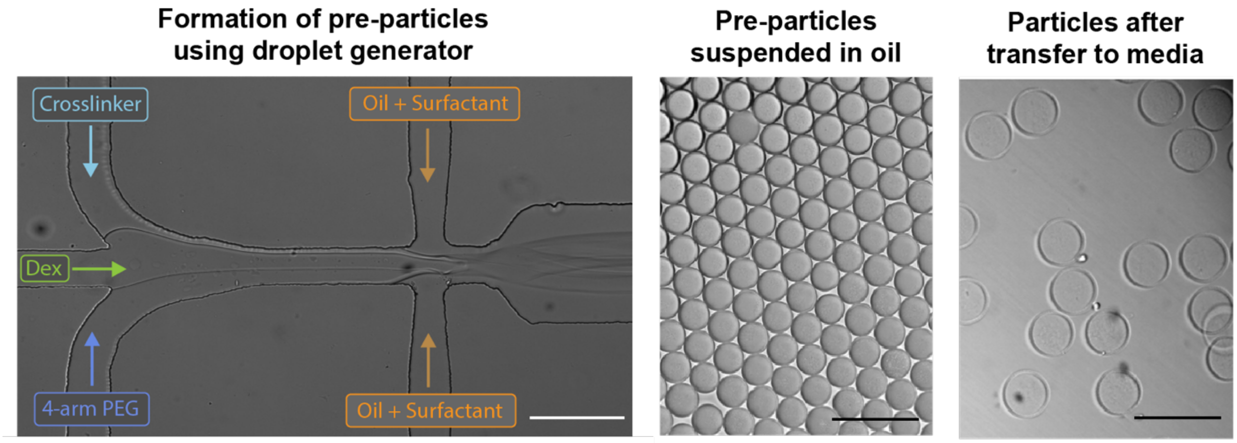


**Figure S1.** Microfluidic fabrication of PicoShells. In the first part of PicoShell fabrication, dextran, 4-arm PEG, and crosslinker are co-flowed immediately before droplet formation in a microfluidic device to form pre-particles. Phase separation and cross-linking occurs in these pre-particles over a 2h period after droplet formation. After the 2h incubation period, the cross-linked particles are transferred from oil to aqueous solution where dextran leaks out of the inner cavity of the particles to form PicoShells. The outer diameter of PicoShells is larger than that of pre-particles (~90 µm and ~70 µm respectively) due to the expected swelling of the PEG hydrogel in aqueous solutions. Scale bars = 200 µm.

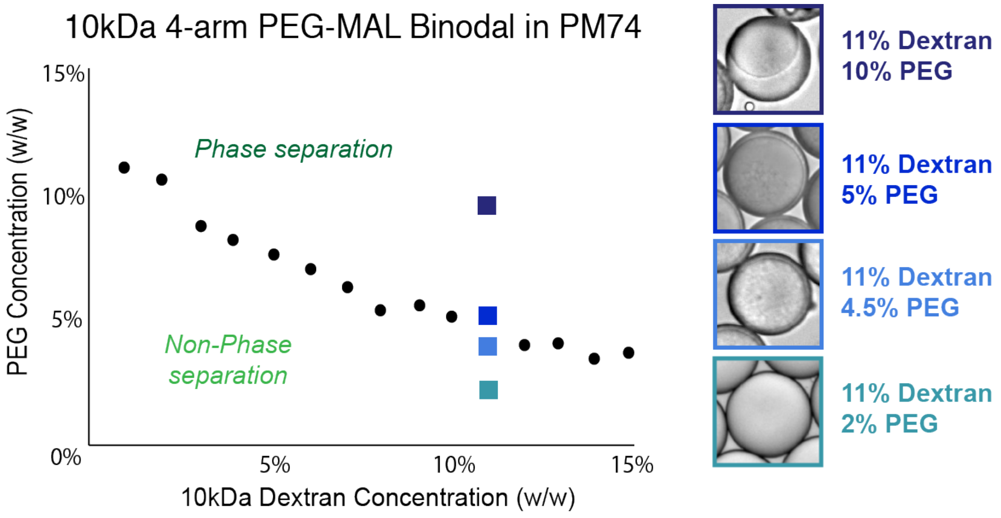


**Figure S2.** Phase diagram of 10kDa 4-arm PEG-maleimide and 10 kDa dextran in PM74 seawater medium. Black dots represent concentrations we experimentally determined where the binodal is located through visual observation of droplets with varying concentrations of PEG and Dextran. PEG and dextran concentrations below the binodal curve do not result in phase separation. PEG and dextran concentrations above the curve result in noticeable phase separation. PicoShells are formed by mixing PEG and dextran at in-droplet concentrations within 0.5-2% above and to the right of the points on the binodal curve.

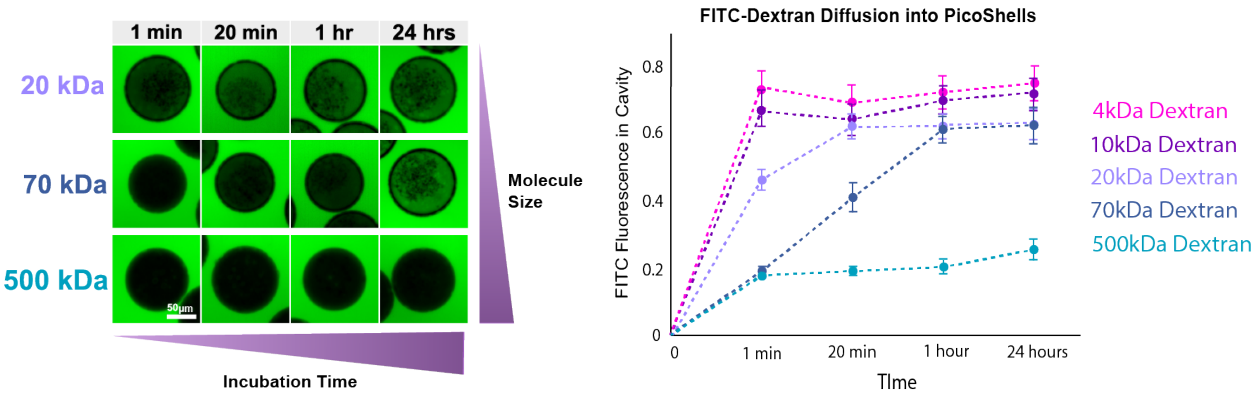


**Figure S3.** Characterization of diffusion through the outer hydrogel matrix of the Picoshells. The rate at which molecules diffuse through the PicoShell's outer matrix depends on the size of the molecules. PicoShells were placed into solutions containing FITC-dextran (green) of various molecular weights and the transport of those molecules into the PicoShell cavity was tracked over time. For FITC-dextran ≤ 20kDa we observe rapid entry into the cavity (i.e. equilibrium within 20min). 70kDa FITC-dextran diffuses more slowly into the cavity, reaching equilibrium at between 1 and 24h. 500kDa FITC-dextran was not seen to enter into the PicoShell cavity even after 24h. Scale bar = 50 μm.

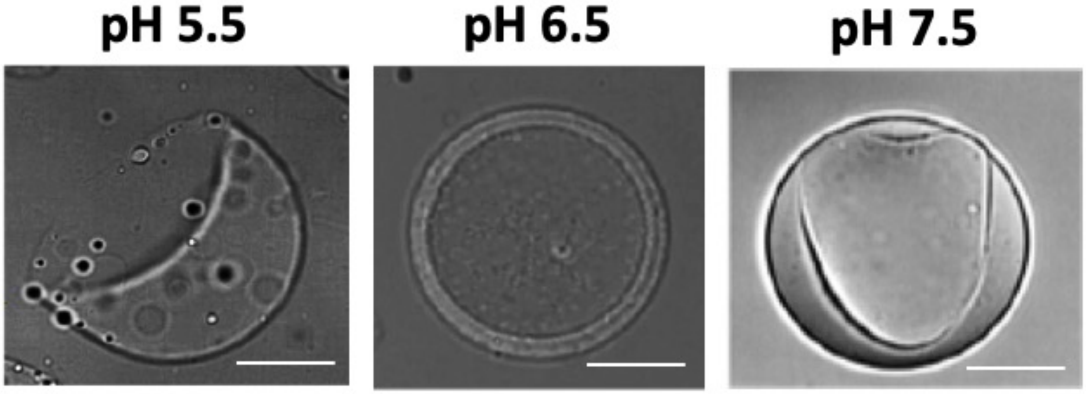


**Figure S4.** Sensitivity of particle shape to crosslinking speed. The rate of crosslinking was varied by adjusting the pH of the precursor solution from 5.5 – 7.5 for a fixed PEG and dextran concentration. At a pH of 7.5, the time between droplet formation and crosslinking occurs too quickly, resulting in polymerization before the PEG and dextran are able to phase separate completely and form a uniform hollow shell structure. At a pH of 5.5, crosslinking rate is reduced resulting in formation of janus particles. We theorize that in the initial phase of crosslinking the effective molecular weight of the PEG monomers increases as they link together leading to a shift in the binodal which changes the equilibrium morphology. At a pH of 6.5 we found that the crosslinking rate is slow enough to alow for complete phase separation and quick enough to preserve the ideal PicoShell morphology. , Scale bar = 25μm.

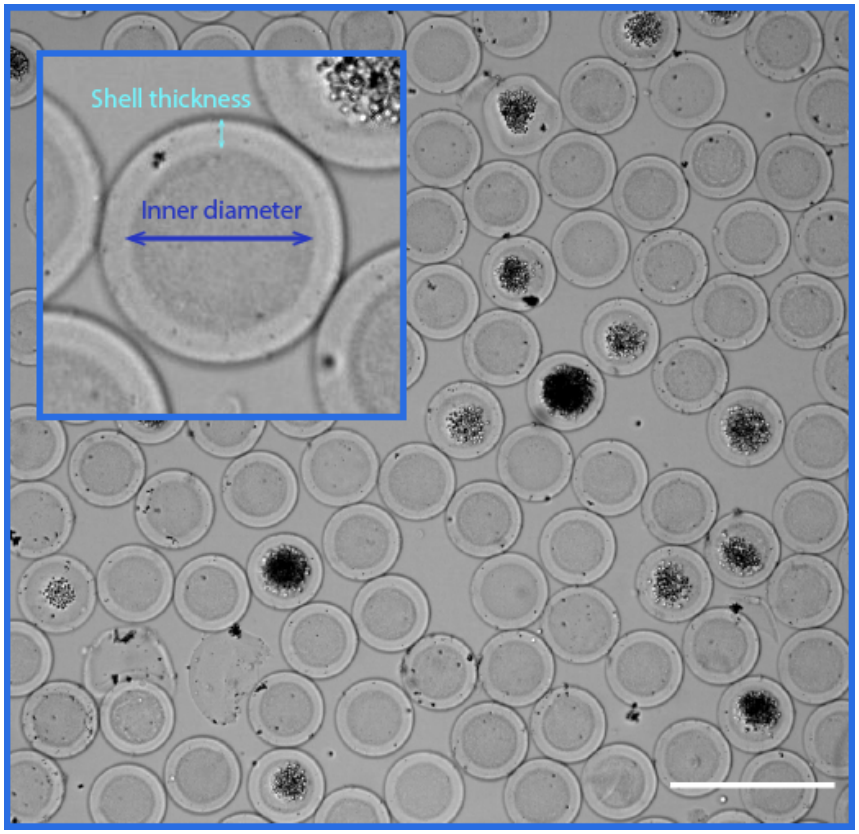


**Figure S5.** PicoShells fabricated with encapsulated *Chlorella* colonies are highly uniform in size and morphology. PicoShells have an average outer diameter of 90.7μm with a CV of 1.7%. The particles also have an average shell thickness of 12.7μm with a CV of 6.9%. Scale bar = 200μm

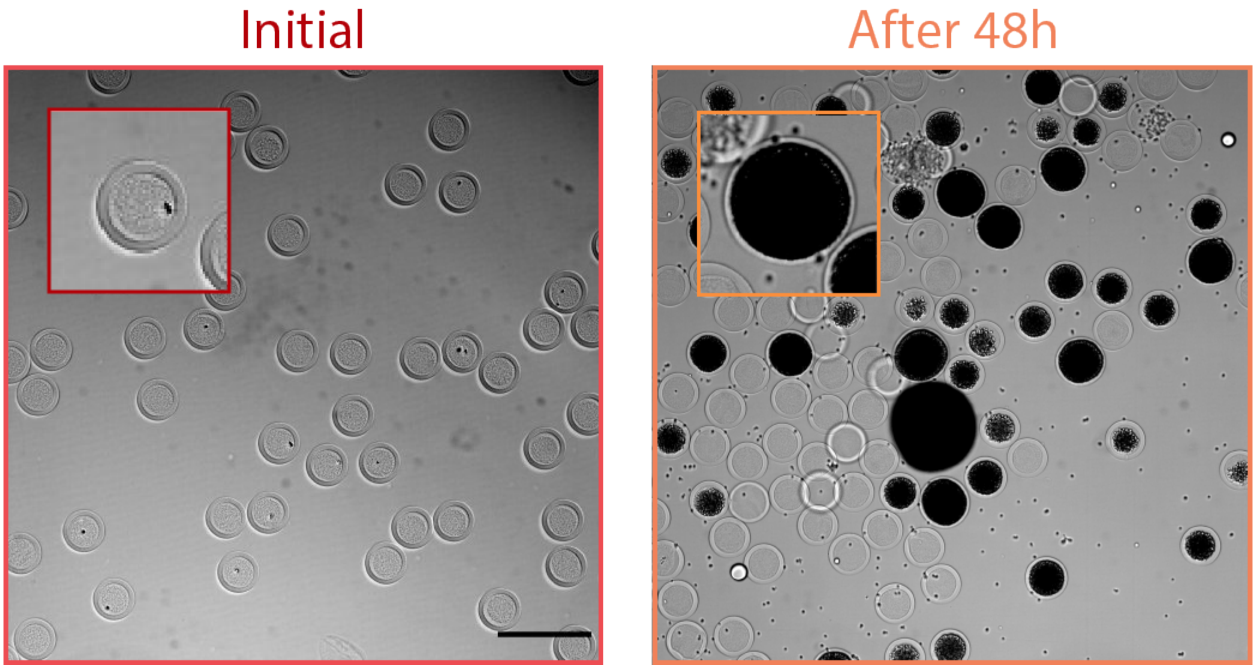


**Figure S6.** Expansion of PicoShells due to *S. cerevisiae* growth. PicoShells containing S. cerevisiae initially have the same outer diameter. However, *S. cerevisiae* can cause PicoShells to stretch and increase in diameter (97µm OD, 11.2% CV at 48h) as the yeast cells multiply over time. Scale bar = 200µm.

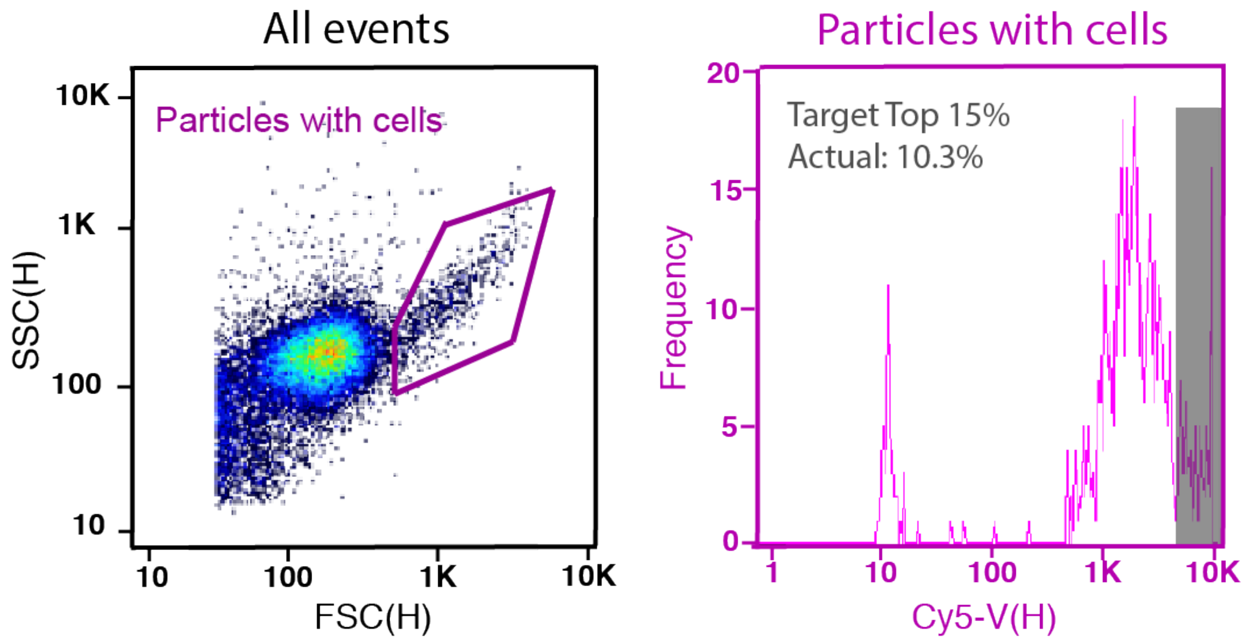


**Figure S7.** Full selection FACS data. For the full selection of hyperperforming *Chlorella* colonies, we first gated events that produced high scatter signal (purple gate) and sorted events that produced the highest 15% [Cy5-V(H)] signal. The data presented only shows 14,854 of the 121,213 PicoShells that were screened for the selection.

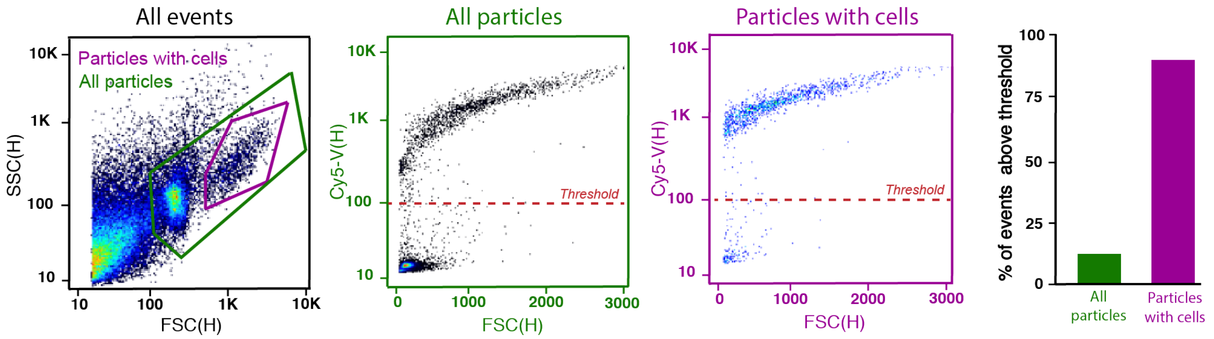


**Figure S8.** Additional *Chlorella*-containing PicoShell screening data. PicoShells containing *Chlorella* clonal colonies produce 3 distinct [FSC(H)] vs [SSC(H)] readout populations: one for particles containing colonies, one for empty particles, and one for debris. When all particles are gated, there are two distinct populations on a [FSC(H)] vs [Cy5-V(H)] plot, one population with a low [Cy5-V(H)] readout and another population with higher [Cy5-V(H)] readouts. When we gate the high scatter readouts from the [FSC(H)] vs [SSC(H)] plot, we find that this population corresponds to PicoShells that produce the highest [Cy5-V(H)] readouts. When all particles are gated, 14.1% of events are above a 100 [Cy5-V(H)] readout. When the high scatter population is gated, 89.9% of events are above a 100 [Cy5-V(H)] readout. Since we anticipate that colony-containing PicoShells will produce high scatter signal (as observed in an inverted microscope) and high Cy5 fluorescence signal due to a *Chlorella's* chlorophyll autofluorescence, we infer that this high scatter, high Cy5 population corresponds to colony-containing PicoShells. When we gate for 'Particles with cells' on the [FSC(H)] vs [SSC(H)] plot, we obtain a 94.0% purity and 72.7% yield of colony-containing PicoShells.

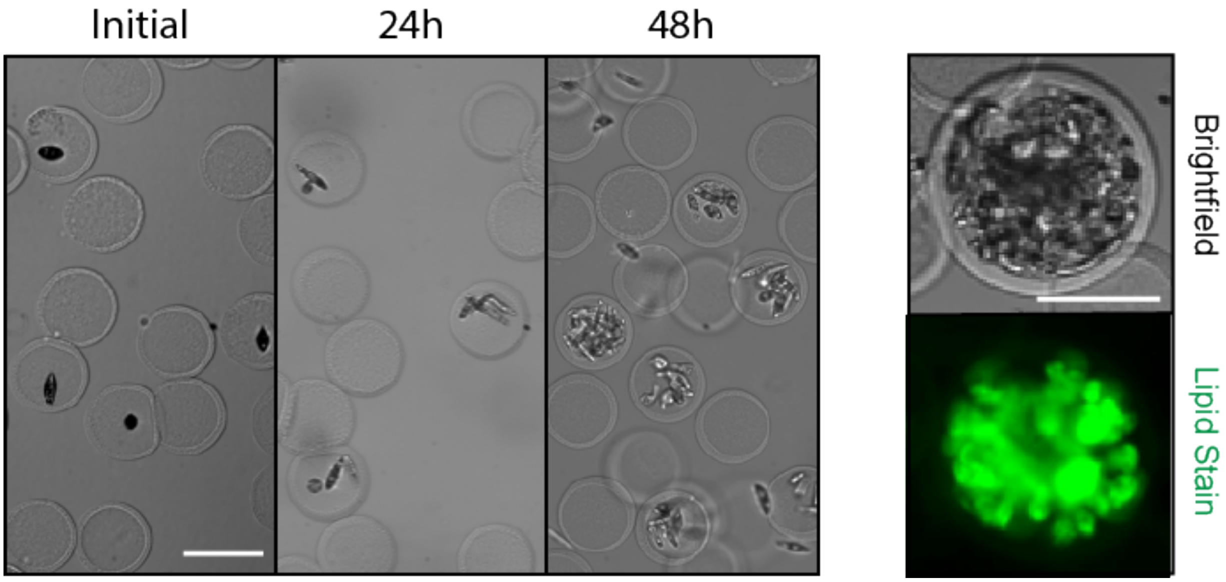


**Figure S9.** *Euglena gracilis* biomass and lipid accumulation within hollow shell particles. (a) Euglena cells were encapsulated into non-degradable hollow shell particles and allowed to grow over a 2-day period. We demonstrated that Euglena are viable and can accumulate biomass within hollow shell particles over this period. Particles were exchanged into a solution with BODIPY after the 2-day incubation to fluorescently label wax esters within the encapsulated microalgae. Stains were able to transport through the solid polymer matrix of the hollow particles and label cells without sticking a significant amount to the particle's PEG surface. Scale bars = 100μm.

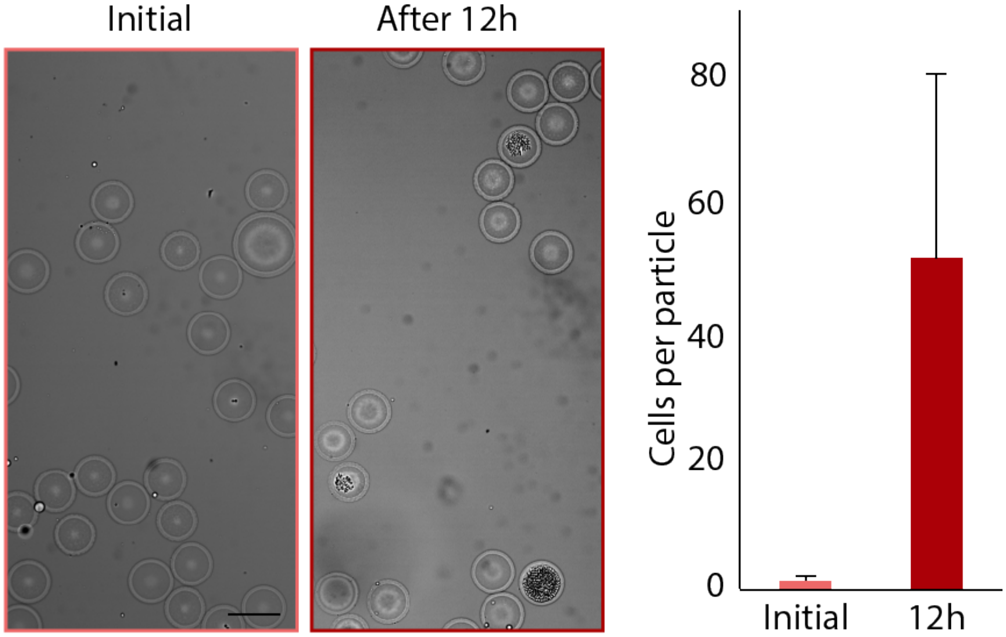


**Figure S10.** Growth of yeast in di-sulfide cross-linked particles. *S. cerevisiae* were encapsulated into PicoShells crosslinked with 20kDa PEG-OPSS and DTT, such that the particle's outer shell contains chemically degradable di-sulfide linkages. Yeast encapsulated into these particles were able to remain viable and multiply, increasing from an average 1.5 cells/PicoShell to 51.4 cell/PicoShell in 12h. Scale bar = 100μm.

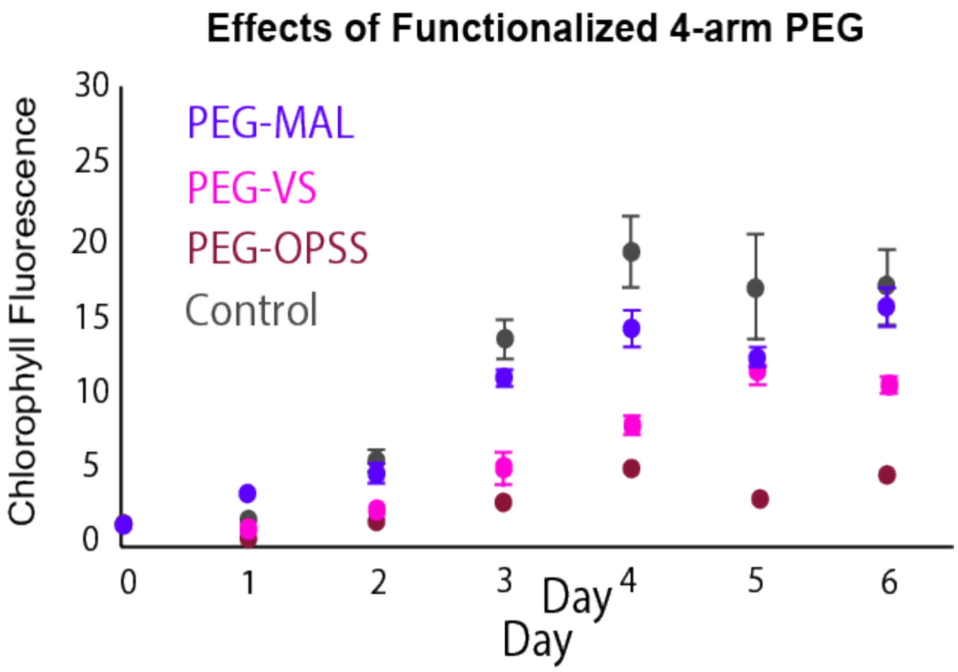


**Figure S11.** Effects of functionalized 4-arm PEG on *Chlorella* viability. *Chlorella* were incubated with 5% (w/w) 10kDa 4-arm PEG-malemide (MAL), 10kDa 4-arm PEG-vinyl sulfone (VS), and 10kDa 4-arm PEG-Ortho-Pyridyldisulfide (OPSS) dissolved in PM74 medium for 2 hours before being transferred into fresh PM74 medium for tracking of biomass accumulation. 4-arm PEG MAL has slight effects on the growth ($P < 0.05$ at time = 3 days) but 4-arm PEG-VS and 4-arm PEG-OPSS has significant effects ($P < 0.001$) the overall of growth of *Chlorella.*

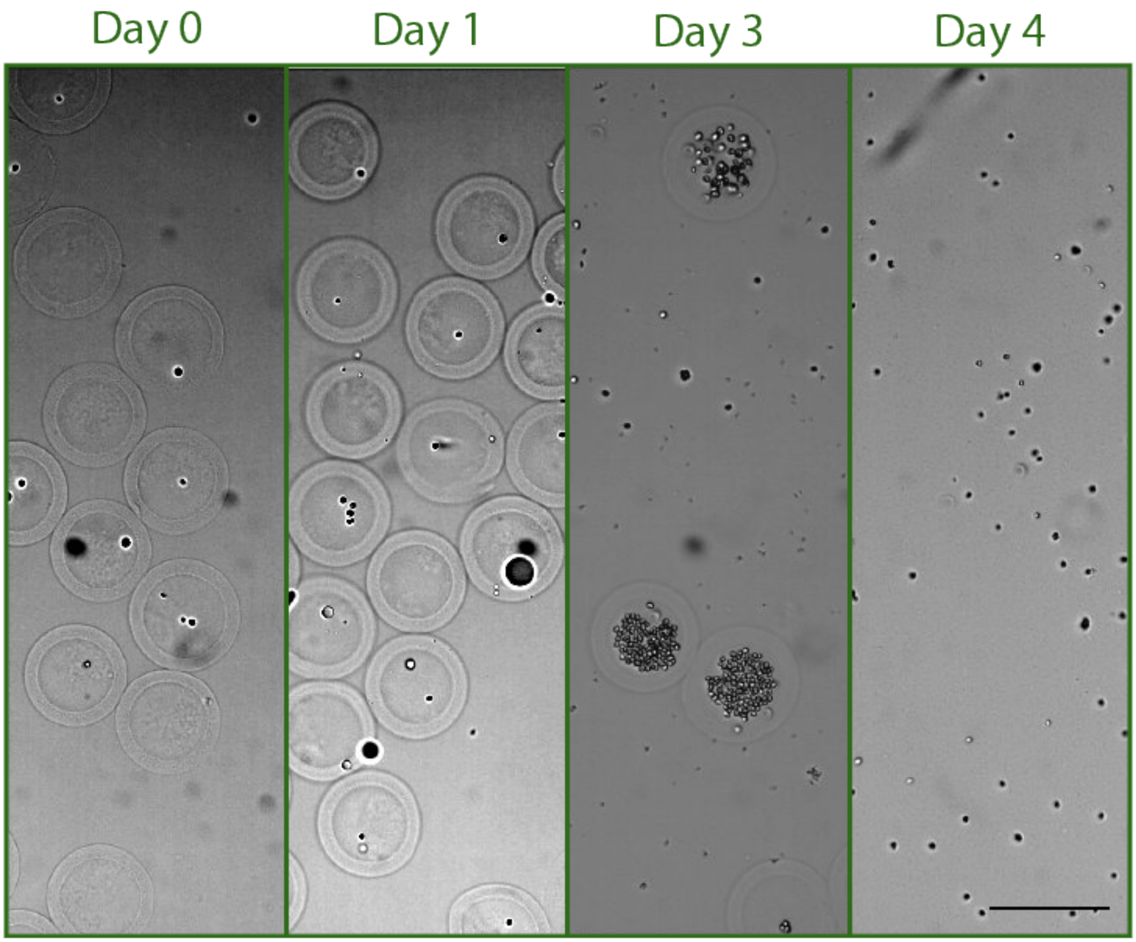


**Figure S12.** Growth of *Chlamydomonas reinhardtii* in MMP-degradable shells. *C. reinhardtii* are capable of growing in PicoShells crosslinked with MMP-degradable peptide crosslinker. However, cells pre-maturely induce the degradation of the PicoShells over time. This is likely due to natural MMPs secreted by *C. reinhardtii*. Scale bar = 100μm.

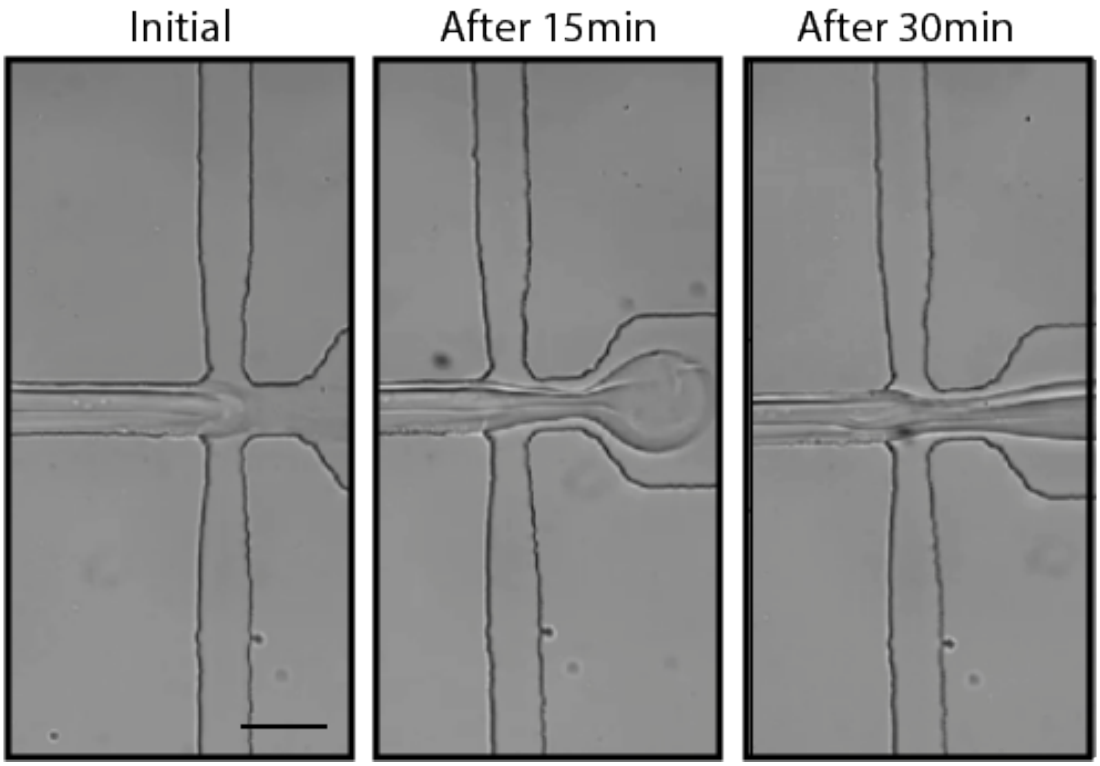


**Figure S13.** Disruption of PicoShell generation over time. Crosslinked material tends to form in the droplet generation junction over time, causing droplet formation to be disrupted. At a pH of 6.5, this usually starts to occur ~15 min after droplet generation is initiated when oversized droplets start to form. After ~30min, no droplets form. The time between initial mixing of reagents and jetting shortens as the pH of the solutions is increased. Scale bar = 100μm.
